# Vernalization regulatory network identifies potential novel functions for *ZCCT* genes within *HvVRN2*

**DOI:** 10.1101/2025.04.29.651237

**Authors:** Francesc Montardit-Tarda, Irene Puyó, Bruno Contreras-Moreira, Ildikó Karsai, Philippa Borrill, Ana M Casas, Ernesto Igartua

## Abstract

- Flowering is a key yield determinant, controlled by environmental and genetic factors. Vernalization requirement, a prolonged cold period to promote flowering, is determined in barley (*Hordeum vulgare* L.) by the allele in *HvVRN1* and the presence of locus *HvVRN2*, which consists of two *ZCCT* genes: *HvVRN2a* and *HvVRN2b*. While the effect of *HvVRN2* presence is well-known, its outcome on the transcriptome and other genes outside the flowering/vernalization pathway remains unknown.
- Near-isogenic lines with presence/absence of *HvVRN2*, were subjected to different lengths of vernalization treatments, identifying the expression dynamics of both *HvVRN2* genes and wider transcriptional responses controlled by VRN2.
- Our work revealed previously unknown differences in expression levels of the two *HvVRN2* genes, in responses to its main repressor, *HvVRN1*, and an effect on tillering. Besides, we shed new light on the repressing mechanism of VRN2 on flowering, using protein modelling. A regulatory network analysis pointed at new candidate genes for flowering regulation related to the vernalization pathway, and suggested additional biological roles for *HvVRN2b*.
- This study re-establishes *HvVRN2* as a central gene for understanding barley responses to environmental cues, beyond its previously accepted simple role, and expands the catalogue of genes related to the vernalization pathway, which may be targeted in barley breeding.

## Introduction

The transition to flowering is a major physiological event in plants. Its timing has a large agronomic impact in cereal crops of the *Pooideae* family (Carrera et al 2024). This process is highly controlled by environmental conditions, and both gene diversity and regulation. Vernalization is the induction of flowering after exposure to a prolonged cold period. This period is necessary in regions with cold winters, in which flowering too early will expose the reproductive tissues to frost risk. During reproductive development, the frost-sensitive floral structures within the elongating tillers are highly exposed to ambient temperatures. Hence, the need to have a built-in genetic mechanism to delay this phase until conditions are favourable. Barley and wheat varieties that require vernalization are named winter growth habit genotypes (or “winter” for short).

The vernalization response is elicited by an appropriate assortment of alleles at certain genes. Three main vernalization-related genes have been identified in barley: *HvVRN1*, *HvVRN2* and *HvVRN3*. The gene *HvVRN1*, or *HvBM5*, is located on chromosome 5H, and encodes for a MADS-box transcription factor which induces flowering (Trevaskis et al 2003, Yan et al 2003), binding directly to the promoter region of *HvVRN3* (Deng et al 2015). The expression of *HvVRN1* in winter genotypes is activated by enough vernalization time (Trevaskis et al 2006). Two paralogous genes of *HvVRN1*, *HvBM3* and *HvBM8*, are found in most cereals and are considered as redundant flowering regulating genes (Li et al 2019). *HvVRN2*, located on chromosome 4H, comprises two genes duplicated in tandem, with Zinc-finger and CONSTANS-like domains (ZCCT) (Yan et al 2004; Dubcovsky et al 2005). A third truncated gene is included in the locus, but is not expressed (Trevaskis et al 2006). *HvVRN2* is expressed in leaves during long days, inhibiting *HvVRN3*. After vernalization, *HvVRN1* is activated, which then represses *HvVRN2* in the leaves (Sasani et al 2009), allowing long-day (LD) induction of *HvVRN3*. The known allelic variation of *HvVRN2* in barley is of presence/absence nature, determining the growth habit of the genotypes: winter (presence) and spring (absence). Moreover, presence of *HvVRN2* affects agronomic performance (Karsai et al 2006). *HvVRN3*, or *HvFT1*, is the ortholog of *Arabidopsis thaliana FLOWERING LOCUS T* (*FT*), promoting the transition to flowering when induced in leaves (Yan et al 2006). *HvFT1* is not expressed in apices (Digel et al 2015). Rather, the protein is transported from the leaves to the apex through the phloem, as evidenced in *Arabidopsis* (Corbesier et al 2007) and rice (Tamaki et al 2007), inducing the reproductive development of the apex due to the florigen signal. Differences in vernalization requirements are mostly modulated by the allelic diversity at these three main vernalization genes, as reviewed in Fernández-Calleja et al (2021).

Even though not considered as classic vernalization genes, CONSTANS-like domain containing genes, *HvCO1* and *HvCO2*, have been proposed to interact with *HvVRN2. HvCO2* is an activator of flowering under LD photoperiod. (Mulki and von Korff 2016). In *Arabidopsis*, CO protein is stable at the end of the day under LD photoperiod, while it is degraded during the morning and night (Brambilla and Fornara 2017). The CO protein interacts with the dimer formed by Nuclear Factor-Y (NF-Y) subunits NF-YB and NF-YC, and the resulting complex recognises specifically the CCACA DNA motif, through the CONTANS-like (CCT) domain (Gnesutta et al 2017). Several CO/NF-YB/NF-YC complexes bind to the various CCACA motifs located in the promoter region of *FT*, interacting among them by the two B-box zinc finger domains of the CO protein, activating *FT* expression (Huang et al 2025). Allelic variation at another photoperiod gene, *HvPRR37* (*PPD*-*H1*) has been found responsible for altered expression of *HvVRN2* (Mulki and von Korff 2016). The genes, *HvCO2*, both *HvVRN2* and *HvPRR37*, present one CCT domain in their protein structure while showing different number of zinc-finger domains: two B-box zinc fingers, one zinc finger and none, respectively. Interactions between these genes and NF-Y subunits were identified in wheat, using yeast two– and three-hybrid assays. VRN2 protein competes with CO2 for interactions with the same NF-Y subunits (Li et al 2011). This competition, and the known roles of VRN2 as repressor and CO2 as activator, suggests an antagonistic role of the two types of complexes on progress towards flowering. Li et al (2011) also found that both CO2 and VRN2 could dimerize, as homo and heterodimers, but their effect on *FT1* expression is unknown. PPD1 can dimerize with CO1 and CO2 proteins, but it does not interact with most of the NF-Y subunits (Shaw et al 2020). The potential role of the CCT protein dimers is not evidenced.

Transcriptome analysis has been used in barley since quite early (Zhang et al 2004), but wide availability of cheap RNA sequencing allowed scaling up the studies, aiming at different plant tissues or conditions (Cantalapiedra et al 2017, Müller et al 2020). Large-scale experiments have been conducted to identify transcriptomic differences or landscapes (Kovacik et al 2024, Thiel et al 2021). The availability of transcriptomic data allowed creating expression databases referenced to the barley reference cultivar Morex (Mascher et al 2021), as BarleyExpDB (Li et al 2023), or for the barley reference transcriptome BaRTv1.0 (Rapazote-Flores et al 2019), as EoRNA (Milne et al 2021). Therefore, most studies are referenced to the genome of Morex, a spring cultivar lacking the gene models of *HvVRN2*. There is abundant information on the physiological effects of *HvVRN2* on the vernalization mechanism. However, there are still knowledge gaps about gene regulatory networks (GRN) inducing flowering in barley, particularly on the involvement of *HvVRN2*. Genes related to *HvVRN2* could be potential targets for barley breeding to fine tune flowering, and for better agronomic performance. We conducted a multifactorial experiment with *HvVRN2* near-isogenic lines and growth conditions chosen to capture the full range of *HvVRN2* expression dynamics. The transcriptome of multiple samples was analysed to reveal the gene regulatory network involving *HvVRN2*. Protein modelling of CCT genes with NF-Y subunits was conducted to suggest potential competition of the CCT/NF-Y complexes and its relevance for flowering regulation.

## Materials and methods

### Plant material, experimental design and tissue harvest

Two BC_4_-near-isogenic lines (NILs) provided by Dr. Ben Trevaskis (CSIRO, Australia) were used in this study. The two NILs, CSIRO01 (C01) and CSIRO03 (C03), differed in the presence/absence of the vernalization gene *HvVRN2*, respectively, while sharing the same alleles at *HvVRN1*, *HvPRR37 (PPD-H1)* and *HvPHYC.* Both genotypes share 97.65% of the genomic background, identified by the 50K Illumina SNP chip (Ochagavía et al 2022). C01 is a true winter genotype (winter allele at *HvVRN1*, presence of *HvVRN2*), whereas C03 is a facultative genotype (winter allele at *HvVRN1*, absence of *HvVRN2*).

Seeds were sown in 5×7 seed trays of 200 cm^3^ each cell (115 mm height, 50×50 mm at the top and 31×31mm at the bottom), in a soil mix of 47% peat moss, 39% sand and 14% vermiculite, with 0.004% of controlled-release fertilizer (Plantacote® Plus 4M; 14-9-15 N-P-K plus 2 Mg_2_O and trace elements). After 3-4 days for seedling emergence, the trays were moved to a vernalization chamber at 4 ± 2 °C with short-day photoperiod of 8/16 hours of light/dark, respectively. Plants were vernalized for either 0, 7, 15, 30 or 60 days, and then moved to a growth chamber under long-day photoperiod, 16/8 hours of light/dark periods, with a temperature of 18/16 °C day/night, a relative humidity of 65% and light intensity of 300 μmol m^−2^ s ^−1^ PAR. The seed tray of 0 days of vernalization, without any vernalization treatment, was directly placed in the growth chamber after seedling emergence. After 5 and 15 days in controlled environment, the basal part of the last-fully expanded leaf of 7 individual plants was sampled at 12 hours into the light period, and immediately frozen in liquid nitrogen. For genotype C01, with presence of *HvVRN2*, another sample was taken after 25 days. Samples were stored at –80 °C until RNA extraction.

Each 5×7 seed tray corresponded to a vernalization time. Three of the five rows were sown with C01, and each one was completely sampled after 5, 15 or 25 days in the growth chamber. The remaining two rows were sown with C03, and sampled at 5 and 15 days. A summary of the experimental design is shown in Fig. 1.

**Figure 1.**
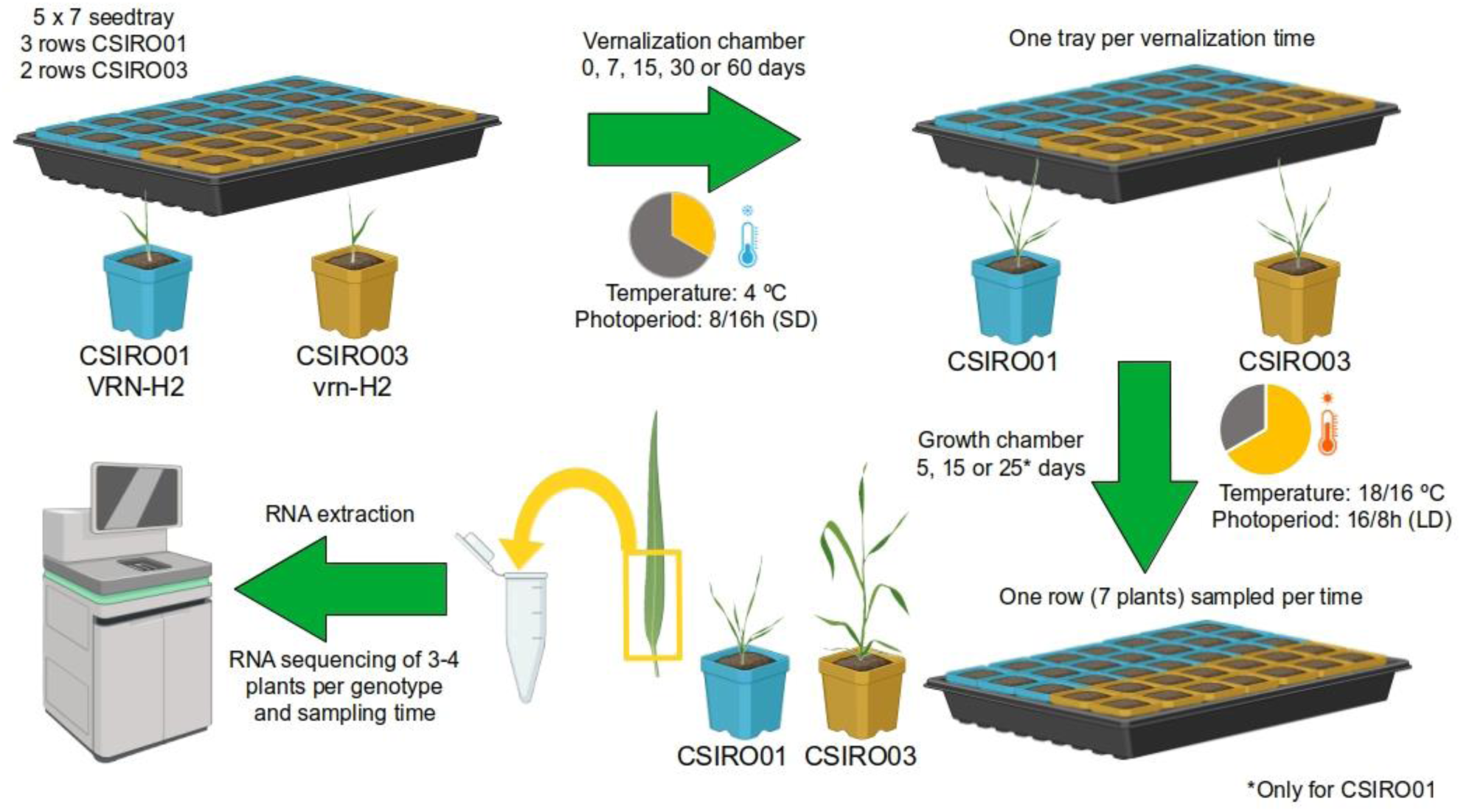
Scheme of the experimental design. Seeds of Near-Isogenic Lines (NILs) differing in presence/absence of *HvVRN2*, CSIRO01 with *HvVRN2* and CSIRO03 without *HvVRN2* were sown in seed trays. After seedling emergence, trays were moved to a vernalization chamber at 4 °C and short-day photoperiod for 7, 15, 30 or 60 days. After the vernalization treatment, plants were grown in standard growing conditions with long-day photoperiod. A tray without vernalization, considered as 0 days of vernalization, was moved to the growth chamber after seedling emergence. The basal part of the last-fully expanded leaf of 7 individual plants per NIL was sampled at 5 and 15 days, and at 25 days for CSIRO01 due to its slower development. RNA was extracted for all leaf samples, and between 2 to 4 were sequenced per genotype, vernalization treatment and sampling timepoint.

Phenotypic data on apex development and tiller number for the two genotypes was measured in the Phytotron facilities at Martonvásar (Hungary), as reported in Ochagavía et al (2022). Four plants per genotype were vernalized at 4 °C for 60 days and then moved to the Phytotron, in long-day photoperiod of 16/8 hours of day/night at constant temperature of 18 °C. Another four plants per genotype were grown in the Phytotron without any vernalization treatment. Tiller number was counted every 3 to 5 days until plants reached stem elongation, stage Z31 (Zadoks et al 1974). Tiller number was counted again when aws were visible, stage Z49. Since unvernalized C01 genotype did not reach Z31, tillers were counted until tiller number was stable in three consecutive measures (day 54). Statistical differences between vernalized genotypes at Z31 and Z49 were analysed with a t-test. Linear models were fitted per genotype and vernalization treatment. Statistical analyses were computed in *R* software v4.4.2 (R Core Team 2023).

### RNA extraction, sequencing and quality control

RNA was extracted from the leaf samples with the Total RNA Mini Kit for Plants (IBI Scientific), following manufacturer instructions, and quantified using a NanoDrop 2000 (Thermo Fisher Scientific). Quality of the RNA extraction was checked in a 1% agarose gel, and determined with QUBIT RNA IQ. RNA samples were diluted at 50-100 ng/μl, when necessary, and sent to Novogene UK for sequencing. After reception, RNA was quality-checked with Agilent Bioanalyzer. Libraries were prepared with poly-A enrichment, and quantified with QUBIT and real-time PCR. Samples were sequenced by Illumina NovaSeq 6000, sequencing of 2 x 150 paired-end reads, for a total of 15 Gb raw data per sample on average. Raw reads were processed with *fastp* software (Chen et al 2018), removing reads with adapters, with N ≥ 10% or with a QScore of over 50% bases ≤ 5. After sequencing and quality control, 2 to 4 biological replicates remained for each combination of genotype and treatment (Table S1).

### Transcriptome quantification and differentially expressed genes

Clean sequence data was pseudo-aligned with *kallisto* software v0.46.1 (Bray et al 2016) to BaRTv2 reference transcriptome (Coulter et al 2022), of the spring barley cultivar Barke. As Barke lacks *HvVRN2*, the transcripts encoded by the two gene models of *HvVRN2* (Horvu_13942_4H01G516500.1 and Horvu_13942_4H01G516600.1, identified as *HvVRN2a* and *HvVRN2b,* respectively) were taken from barley pangenome v1 (Jayakodi et al 2020), specifically from the Spanish winter landrace HOR_13942 (ERS4201453), and added to the reference transcriptome. Transcript-level estimated counts were normalized into transcripts per million (tpm) (Table S2), and then aggregated to gene-level with *tximport* package v1.34.0 (Soneson et al 2015) in *R* software v4.4.2. Uncertainty of the estimated expression was considered with 100 bootstraps.

Principal component analysis (PCA) and identification of differentially expressed genes (DEGs) were carried out with *sleuth* package v0.30.1 (Pimentel et al 2017). Genes considered for analysis required at least 5 estimated counts in 25% of the samples. DEGs were identified with a likelihood ratio test comparing the following two models, aiming to pinpoint DEGs specifically influenced by vernalization:

> *reduced model: days under long-day photoperiod*
>
> *full model: days under long-day photoperiod + vernalization*

The analysis was carried out for the two genotypes separately. Genes were considered as differentially expressed by a multiple test correction adjusted p-val (q-val ≤ 0.01).

### Identification of co-expressed genes

Clustering in modules of co-expression was performed at gene-level by *WGCNA* package v1.73 (Langfelder and Horvath 2008), which uses a correlation network approach. Genes were selected if they presented 10 counts in at least 9 samples. Gene expression in counts was variance-stabilizing transformed with *DESeq2* package v1.46.0 using the parameter blind = TRUE (Love et al 2014). Gene and samples outliers were checked, and filtered out if necessary. A soft-power threshold was calculated as the first power to exceed a scale-free topology fit index of 0.9 (β = 4), considering a signed hybrid correlation network and biweight midcorrelations. Parameters for network construction were selected by comparing different combinations, considering the total number of identified modules and flowering-related genes membership. The topological overlap matrix (TOM) was created with an unsigned TOMtype approach. The minimum number of genes per module was 40 (minSize= 40). Dendrogram was cut at 0.97 for module detection (detectCutHeight = 0.97) but module splitting was set to mid-sensitive (deepSplit = 2) and module merging was allowed (mergeCutHeight = 0.15). Module eigengene (KME) was calculated for all genes and modules. Modules were correlated to the factors of the experimental design.

### Gene Ontology enrichment

Modules of co-expression were enriched by Gene Ontology (GO) with *clusterProfiler* package v4.16.6 (Wu et al 2021). The GO term annotation of BaRTv2 was complemented with the GO terms of both *HvVRN2* gene models from the HOR_13942 genome annotation.

### Transcription factor annotation

For further analysis, identification of transcription factor (TF) gene models of BaRTv2, including both *HvVRN2* gene models from HOR_13942, was required. Protein sequences of all transcripts were analyzed with the *iTAK* Web server v18.12 (Zheng et al 2016). Annotation at gene-level considered the longest transcript.

### Gene regulatory network construction

A gene-regulatory network was constructed with the expression data, filtered as previously described for co-expression clustering, and a list of transcription factors using a random forest approach, with *GENIE3* package v1.28.0 (Huyn-Thu et al 2010). Relationships between genes were filtered with an edge weight ≥ 0.01. Additionally, the network was filtered by known flowering-related genes (Table S3), to focus on the pathways related to flowering and vernalization, and for interactions between transcription factors Network was constructed with *GGally* package v2.2.1 (Schloerke et al 2024) using the Fruchterman-Reingold algorithm, and then manually modified with *Cytoscape* v3.10.3 tool (Shannon et al 2003).

### Pangenome homology of expressed genes

Collinearity-based gene homology between the reference transcriptome (BaRTv2) and barley pangenome v1 was identified with *GET_PANGENES* (Contreras-Moreira et al 2023) and saved as file pangene_matrix_genes.tr.tab (v04102024), available at https://github.com/eead-csic-compbio/barley_pangenes. The resulting pangene set was also used to plot the genomic context of gene *HvSNF2*, linked to *HvVRN2*, with script check_evidence.pl from *GET_PANGENES*.

### Modelization of protein complex structures

Protein complexes of NF-Y domains with other proteins carrying a CCT domain (HvCO1, HvCO2, HvVRN2 and HvPRR37) were modelled with Alphafold 3 Web server (Abramson et al 2024). As a proof-of-concept, HvVRN2a was modelled with other binding partners of Protein Data Bank entries 7C9O and 7CVO, produced for rice (Shen et al 2020) and for Arabidopsis thaliana (Lv et al 2021), obtaining high-scoring complexes. The amino acid sequences encoded by NF-YB and NF-YC genes in barley (Panahi et al 2019), enriched with the annotated high-confidence genes as “Nuclear Factor Y” in the MorexV3 genome (Table S4), were clustered to avoid redundancy with *CD-HIT* v4.8.1 (Li and Godzik 2006), with a sequence identity cutoff of 0.85. A NF-Y protein sequence was selected per cluster (12 for NF-YB and 9 NF-YC), considered as representative. Multiple sequence alignments (MSA) of all representative NF-Y against the NF-YB11 or NF-YC2 proteins of the *Oryza sativa* Japonica group 7C9O complex were performed with *Clustal omega* tool v1.2.4 (Sievers and Higgins 2014). Sequences were trimmed at the start and end according to the MSA, with 7C9O sequences as reference, to reduce noise added by sequences outside the macrocomplex. Models were created for all combinations of the complexes of NF-YB and NF-YC subunits with HvCO1, HvCO2, HvVRN2a or HvPRR37, and an oligonucleotide of length 31 with the CCACA motif extracted from the promoter region of *HvVRN2a* of HOR_13942 (Table S4). The stability of the modelled complexes was calculated as the sum of the values of iPTM and PTM metrics, as proposed in Homma et al (2023).

The position of the CCACA motif across the promoter region (–1000 bp, 0 bp) of the genes coding CCT domain proteins were scanned with *matrix-scan-quick* of the RSAT software (Santana-Garcia et al 2022). Promoter sequences were extracted, using the coordinates available at Table S4. from the genome of HOR_13942 (ERS4201453) from the pangenome v1 (Jayakodi et al 2020).

## Data availability

All sequencing reads are available under study number PRJEB85891 in the European Nucleotide Archive (ENA). Scripts are available at https://github.com/chesQhub/barley_vernalisation_rnaseq.

## Results

### Gene expression changes are mediated by *HvVRN2* and vernalization

To investigate how vernalization affects gene expression through *HvVRN2*, two barley NILS, with *HvVRN2* (C01) or without *HvVRN2* (C03), were exposed to vernalising temperatures for 0, 7, 15, 30 or 60 days before transferring to standard growth conditions for 5, 15 or 25 days (Fig. 1). We hypothesised that these varying conditions would generate quantitative effects on vernalization responses, which could be detected through gene expression changes. Gene expression of the basal part of the youngest fully expanded leaf was measured in each combination of vernalization and growing time per genotype, sequencing mRNA.

Although the focus of the experiment is vernalization, a PCA analysis revealed that plant age under a long-day photoperiod explained more variance than vernalization for both genotypes (Fig. 2A). At the time of sampling, genotypes were in different development stages across treatments (Fig. S1). Moreover, C01 produced more tillers than C03, independently of the vernalization treatment (Fig. S2). DEGs affected by vernalization were identified for each genotype separately. DEGs related to LD growing conditions were discarded using a likelihood ratio test between two fitted models, one with only the LD treatment and another combining the LD treatment with vernalization (Table S5). The intersection of the sets of DEGs between both genotypes marked which genes were affected by vernalization alone or by the effect of both *HvVRN2* presence and vernalization. A total of 769 DEGs were identified only in C01, which were the result of the presence and expression of *HvVRN2* (Fig. 2B). The gene with the lowest q-val in C01 due to the vernalization factor was *HvFT1* (Table 1). The two *HvVRN2* genes were the third and seventh, sorting the DEGs by increasing q-val. Other differentially expressed genes identified include *HvPRR95*, a circadian-clock related gene; a bHLH transcription factor; several zinc fingers, including two orthologs of the rice gene *PROG1* (BaRT2v18chr4HG216300 and BaRT2v18chr4HG216310), which controls plant architecture; a CRT-binding factor (CBF), an AP2/B3 transcription factor or a bZIP transcription factor (BaRT2v18chr5HG259930), ortholog of the rice gene *BZIP77 / FD1*, related to flowering control. (Table 1).

**Figure 2.**
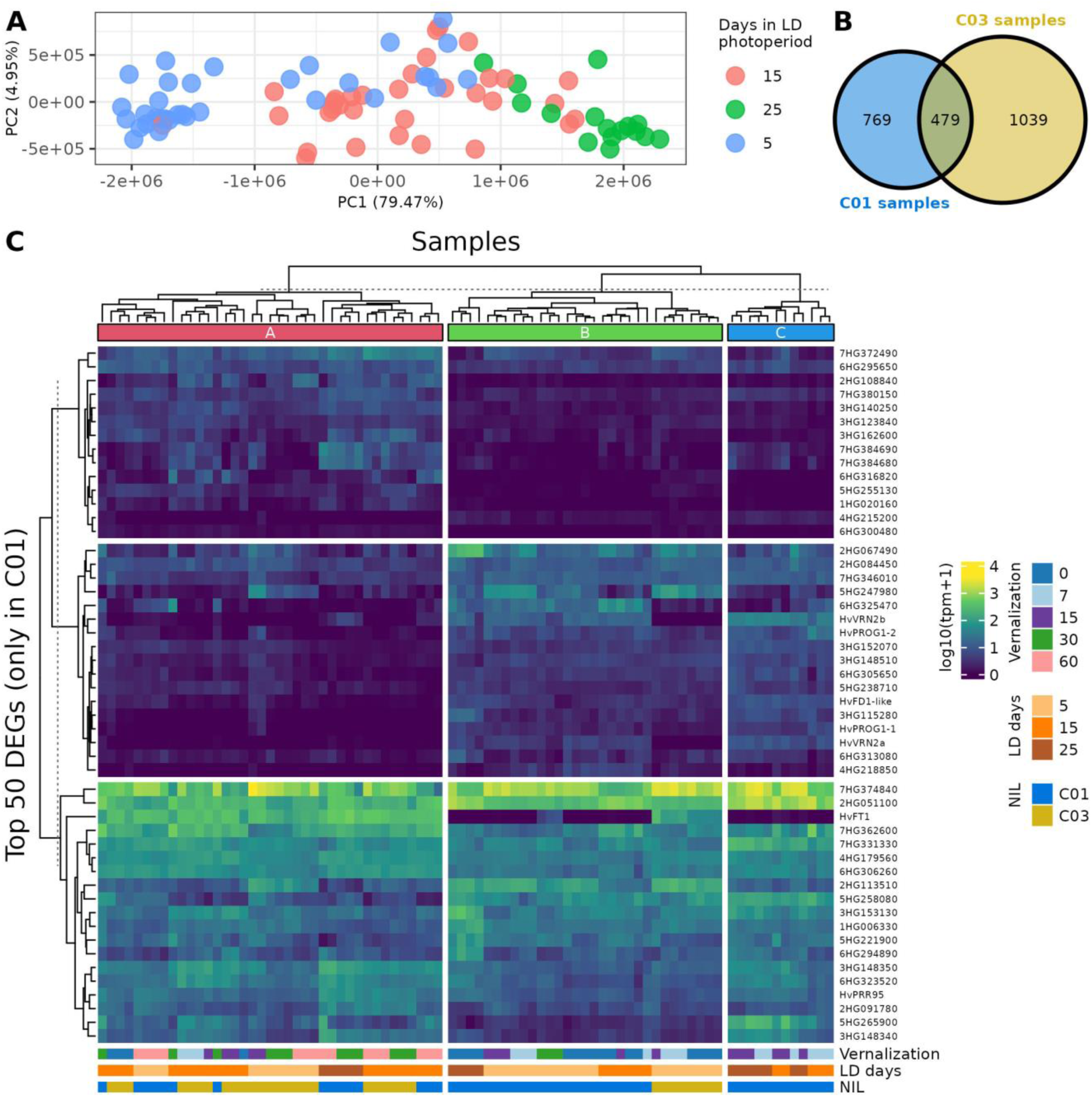
Transcriptome-level differences between samples and differentially expressed genes by vernalization and presence of *HvVRN2*. A) Principal component analysis (PCA) of all samples, coloured by days under long-day photoperiod. B) Venn Diagram of differentially expressed genes (DEGs) detected for each genotype, separately. Blue for C01 samples, with presence of *HvVRN2*, and yellow for C03 samples, without *HvVRN2*. C) Heatmap of the logarithmic expression across all samples of the top 50 DEGs by vernalization, identified only in C01 samples. Samples, in the x-axis, clustered in three major groups. Genes are identified by either their gene name or their BaRTv2 gene model ID, trimming “BaRT2v18chr” to shorten its length, i.e. 3HG148340 as BaRT2v18chr3HG148340.

**Table 1.** Selected differentially expressed genes influenced by vernalization. Column DEG differentiates if the gene was identified only in C01 samples, with presence of *HvVRN2*, or in both genotypes. BaRT2v and MorexV3 ID columns include shortened gene model IDs, trimming “BaRT2v18chr” and “HORVU.MOREX.r3.” from the start of the gene model ID, i.e. *HvFT1* trimmed IDs are 7HG338110 as BaRT2v18chr7HG338110 and 7HG0653910 as HORVU.MOREX.r3.7HG0653910, BaRT2v and MorexV3 respectively.

| Gene | BaRT2v ID | q-val | DEG | MorexV3 ID | MorexV3 annotation |
| --- | --- | --- | --- | --- | --- |
| *HvFT1* | 7HG338110 | 2.01E-62 | C01 | 7HG0653910 | Flowering locus T |
| *HvVRN2a* | Horvu_13942_4H<br>01G516500 | 4.52E-16 | C01 | NA | NA |
| *PROG1-1* | 4HG216300 | 4.92E-13 | C01 | 4HG0411820 | Zinc finger protein, putative |
| *HvVRN2b* | Horvu_13942_4H<br>01G516600 | 1.21E-12 | C01 | NA | NA |
| *bHLH* | 6HG305650 | 2.29E-11 | C01 | 6HG0592420 | Basic helix-loop-helix transcrip-<br>tion factor |
| *HvPRR95* | 5HG258900 | 1.21E-10 | C01 | 5HG0498830 | Pseudo response regulator |
| *PROG1-2* | 4HG216310 | 1.30E-09 | C01 | 4HG0411840 | Zinc finger protein, putative |
| Zinc finger | 3HG115280 | 1.58E-09 | C01 | 3HG0218720 | Zinc finger family protein |
| CBF | 5HG258080 | 1.11E-08 | C01 | 5HG0497560 | CRT-binding factor |
| *AP2/B3* | 4HG215200 | 1.42E-07 | C01 | 4HG0410010 | AP2/B3 transcription factor family<br>protein |
| *HvFD1-like* | 5HG259930 | 5.02E-07 | C01 | 5HG0500770 | Transcription factor |
| *B3* | 4HG206880 | 3.79E-06 | C01 | 4HG0397260 | B3 domain-containing protein |
| *Auxin-respon-*<br>*sive* | 1HG013580 | 5.80E-06 | C01 | 1HG0029890 | Auxin-responsive protein |
| *GATA* | 4HG219040 | 9.56E-03 | C01 | 4HG0416070 | GATA transcription factor |
| *HvVRN1* | 5HG265940 | 2.51E-62 | both | 5HG0511210 | MADS box transcription factor |
| *HvFT2* | 3HG128710 | 1.38E-47 | both | 3HG0244930 | Flowering locus T |
| *HvBM8* | 2HG079040 | 1.10E-44 | both | 2HG0156870 | MADS-box transcription factor |
| *bZIP* | 6HG320060 | 8.30E-25 | both | 6HG0619650 | BZIP transcription factor |
| *HvBM3* | 2HG066580 | 1.75E-21 | both | 2HG0127410 | MADS-box transcription factor |
| *CBF* | 5HG258090 | 1.09E-16 | both | 5HG0497570 | CRT-binding factor |
| *HvPRR37* | 2HG056170 | 2.35E-14 | both | 2HG0107710 | Pseudo-response regulator |
| *CBF* | 5HG258070 | 6.97E-14 | both | 5HG0497540 | CRT-binding factor |
| *VRT2* | 7HG343770 | 1.00E-12 | both | 7HG0664320 | MADS box transcription factor |

In addition, 479 DEGs shared by both genotypes were differentially expressed by the vernalization factor (Table 1, Table S5). Among those, the MADS-box gene *HvVRN1,* and its paralogs (*HvBM3* and *HvBM8)*, an *FT-*like gene (*HvFT2)*, CRT-binding factors, *HvPRR37*, the MADS-box Short Vegetative Protein *HvVRT2* gene or a bZIP (BaRT2v18chr6HG320060).

A clustering of all samples of both genotypes, according to the top 50 DEGs due to vernalization only identified in C01 samples (Table S5) resulted in three major groups (Fig. 2C). The first cluster (A, red) included most of the samples of C03 and C01 from the two longest vernalization treatments, 30 and 60 days. Those vernalization periods were enough to repress the expression of *HvVRN2* in C01 samples, by activation of the expression of *HvVRN1*. As a consequence, *HvFT1* was expressed, triggering the transition to the reproductive phase. The second cluster of samples (B, green) included most of the samples at 5 days for both genotypes, and could be considered as young plants, still unaffected by the long photoperiod conditions of the growth chamber. Unexpectedly, all samples at 0 days of vernalization for C01 genotype were grouped there, independently of the time under long-day photoperiod (5, 15, or 25 days), indicating that they stayed in a similar non-responsive state, as the young plants of other treatments, and did not advance further in their development. The third cluster (C, blue) only grouped C01 samples not fully vernalized (7 and 15 days of vernalization), collected after 15 and 25 days under long photoperiod. In these samples, the transition to reproductive phase was still blocked due to the expression of both *HvVRN2* genes, but their independent clustering indicates that these samples had undergone gene expression changes beyond those observed in plants that experienced no vernalization at all, or that were too young (cluster B). These changes blocked the transition to reproductive phase, and suggest that cluster C plants were in a more advanced vegetative phase, different to young plants of cluster B.

### *HvVRN2* genes are repressed by enough vernalization but activated by partial vernalization

Overall, the expression of the two *HvVRN2* genes followed expectations, according to standing hypotheses, i.e., it is induced by long days and repressed by full vernalization. However, two striking differences were found. Both genes maintained a rather high expression in C01, up to the vernalization treatment of 15 days. With longer vernalization, they were repressed more or less proportionally to the increased expression of their repressor, *HvVRN1*. *HvVRN2b* had much higher expression than *HvVRN2a*. Besides, *HvVRN2a* was completely repressed at some time points, whereas *HvVRN2b* showed some expression across all experimental conditions, even well after *HvVRN1* was fully induced and *HvFT1* was already activated (Fig. 3).

**Figure 3.**
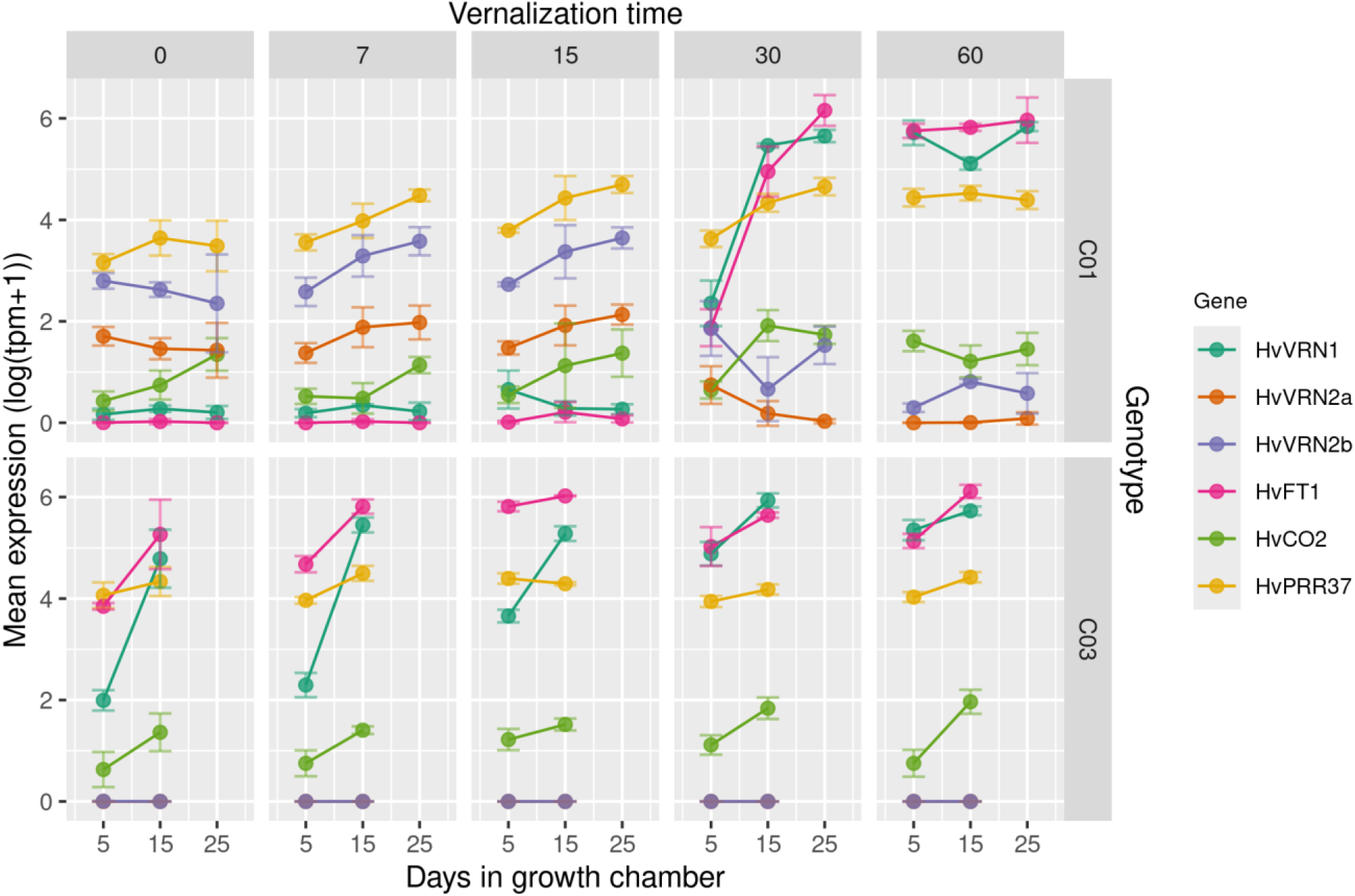
Expression of relevant flowering-related genes in both genotypes. Values are the mean expression of the logarithm of transcripts per million (tpm) + 1. Error bars indicate standard deviation. Horizontal panels differentiate the near-isogenic lines, C01 (top) with presence of *HvVRN2* and C03 (bottom) without *HvVRN2*. Each vertical panel is a vernalization time treatment of 0, 7, 15, 30 or 60 days (left to right). Vernalization panels are divided by each of the sampling points at long-day photoperiod in the growth chamber, at 5 and 15 days. C01 plants were also sampled at 25 days.

Additionally, we observed a certain induction of the two *HvVRN2* genes due to partial vernalization. According to the standing hypothesis, their expression is induced by long days. However, when plants were partially vernalized (7, 15 days of cold treatment), both *HvVRN2a* and *HvVRN2b* increased steadily in expression, in plants that still did not reach jointing stage (Z31). This increase in expression did not occur in plants that were not exposed to cold conditions (0 days vernalization), while *HvVRN2* expression was kept at lower levels after longer vernalization periods (30, 60 days), probably due to the induction of *HvVRN1* expression. This suggests that additional genes influence *HvVRN2* expression.

Gene *HvCO2* was not considered as a DEG, although its q-val in C01 was lower than 0.05 (q-val = 0.029) (Table S5). Its overall expression level across samples was low. This gene showed lower expression than both *HvVRN2* genes in non-fully vernalized samples (0, 7 and 15 days of vernalization). With more vernalization time and with the repression of *HvVRN1*, *HvCO2* expression was higher than that of *HvVRN2*.

### Co-expression clustering identified three vernalization-related gene modules

Co-expressed genes follow similar patterns of activation/repression, as a response to environmental conditions or are regulated by shared TF. We carried out a co-expression clustering to identify which genes are related to the vernalization factor and which are co-expressed with *HvVRN2,* as they could be potential genes related to the flowering/vernalization pathway, as activators or repressors. Genes were clustered in modules by their co-expression (Table S6), with WGCNA, identifying 30 modules. The module size varied between 61 and 2801 genes and included 50% of the total expressed genes. 7558 genes were grouped in an extra module, M0, as not enough co-expression with others was detected.

*HvVRN2b* was included in the co-expression module M29 with another 85 genes, while *HvVRN2a* was placed in the module M0, probably due to its low expression overall. Interestingly, the closest gene to the *HvVRN2* loci on chromosome 4H, the *SNF2 Helicase* (BaRT2v18chr4HG219190), was co-expressed with *HvVRN2b*. A novel MADS-box transcription factor (BaRT2v18chr7HG353720) was also located in this module. *HvVRN1* was located within module M20, same as *HvBM3* and *HvBM8*. Module M20 contained 315 genes, including several *FT* genes (*HvFT1*, *HvFT2* and *HvFT4*), FD-like genes *(HvFD-like 15*), CONSTANS-like genes (*HvCO2*) and genes related to the circadian clock (*HvPRR95* and *HvPRR37*). Another remarkable module was M18, which included genes such as CBFs, *HvVRT2*, and novel identified DEGs such as BaRT2v18chr4HG216300, one of the orthologs of rice gene *OsPROG1*, and BaRT2v18chr5HG259930, ortholog of rice *OsBZIP77 (OsFD1)*.

A correlation between modules and the three experimental factors of design (genotype or presence/absence of *HvVRN2*, time of vernalization and growing days under LD) identified which modules were mostly related to vernalization and presence of *HvVRN2* (Fig. 4A). Modules M18, M20 and M27 were highly correlated to vernalization. M18 and M27 eigengene values decreased during longer vernalization periods while M20 increased (Fig. 4B). M28 and M29 were highly correlated to the genotype background or absence/presence of *HvVRN2* locus, respectively. Clustering the correlation of the modules by the experimental factors identified 7 branches in the dendrogram, differentiating modules M18 and M27, as M18 seems more correlated to C01 genotype while the M27 cluster is to the time in growth chamber (Fig. 4). This suggests an effect of the presence of *HvVRN2* on the expression of the genes within M18.

**Figure 4.**
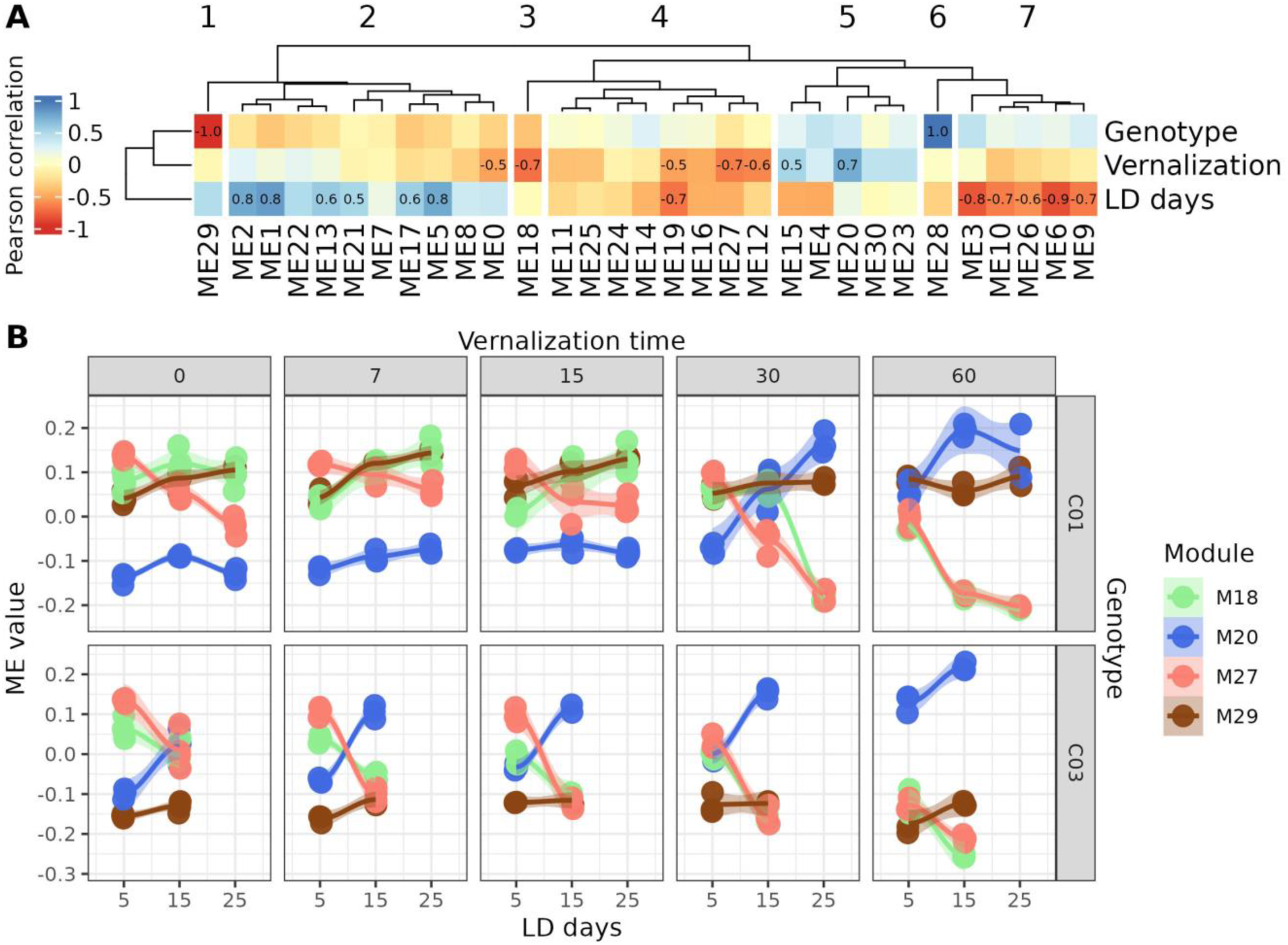
Co-expression modules related to vernalization and presence of *HvVRN2* locus. A) Heatmap of the correlation between co-expression module eigengenes with the factors used in the experimental design, clustering them in 7 groups by dendrogram height. Cell values indicate the correlations above an absolute value of 0.5. The Near-Isogenic Lines, genotype factor, were converted to a binary value as 0 for C01, with presence of *HvVRN2*, and 1 for C03, without *HvVRN2*. B) Eigengenes values, the overall module expression, of the modules related to vernalization (M18, M20 and M27) and presence of *HvVRN2* (M29) throughout all variables of the experimental design: genotype, vernalization time and days of growth under long-day (LD) photoperiod. A smooth line was fitted to visualize differences between samples of each treatment.

Gene Ontology (GO) enrichment of the modules provided some insight on the functions prevalent for each gene cluster (Table 2; Table S7). The GO enrichment of M20 showed terms related to shoot system development and DNA-binding transcription factor activity. *HvVRN1* and *HvFT1* were part of this co-expression module, as other flowering-related genes. Even though not highly correlated to vernalization, M23 was clustered alongside M20 and its enrichment resulted in terms related to chloroplasts and, more importantly, to ovary development. M29, the module with *HvVRN2b*, was enriched for regulation of translational elongation, hinting at a possible mechanism of action for blocking flowering transition. M18 was related to hormonal response while M27 was not enriched by any term.

**Table 2.** Co-expression modules identified by WGCNA, detailing number of genes per module, flowering genes within each module and a summary of the Gene Ontology enrichment.

| Module | Number of genes | Flowering genes | Summarized GO |
| --- | --- | --- | --- |
| M0 | 7558 | *HvVRN2a, HvMADS57, HvFT3, HvCO6, HvBM1* | --- |
| M1 | 2801 | *HvFD-like, HvCO16* | DNA repair |
| M2 | 2778 | *HvNF-YA4* | Ribosome |
| M3 | 2777 | *HvCO3* | Chloroplast, photosystems |
| M4 | 1170 |  | Chloroplast |
| M5 | 1159 | *HvCO18* | Pollen |
| M6 | 898 | *HvPRR73, HvPhyC, HvBM10* | Light response |
| M7 | 793 |  | Chromatin organization, helicase activity, histone modification, vacuole organization |
| M8 | 626 |  | Response to biotic stress |
| M9 | 479 | *HvPRR1* | ADP binding, ADN integration |
| M10 | 451 |  | Cytoskeleton |
| M11 | 416 |  | Cell wall, cell organization |
| M12 | 387 | *HvTOE1, HvCO8* | Cell wall biogenesis |
| M13 | 373 | *HvGI* | mRNA processing, gene silencing by RNA |
| M14 | 368 | *HvCO9* | Oxidative stress, cell adhesion |
| M15 | 362 | *HvPRR59* | NA |
| M16 | 350 |  | ATP, cell wall |
| M17 | 345 |  | Endocytosis |
| M18 | 342 | *HvVRT2, HvFD1* | Response to hormones |
| M19 | 342 |  | Toxin activity, encapsulation |
| M20 | 315 | *HvVRN1, HvBM3, HvBM8, HvFT1, HvFT2, HvFT4, HvFD-H15a, HvCO2, HvPRR37, HvPRR95* | DNA-binding, shoot system development |
| M21 | 302 |  | Response to biotic stress |
| M22 | 295 |  | NA |
| M23 | 276 | *HvFRI-like, HvCMF3* | Stroma, ovule development |
| M24 | 258 |  | Chromatin/histone binding and organization |
| M25 | 245 |  | Nucleosome, epigenetic regulation of gene expression |
| M26 | 204 |  | NA |
| M27 | 167 | *HvVGT1, HvCMF1* | NA |
| M28 | 111 |  | NA |
| M29 | 86 | *HvVRN2b* | Regulation of translational elongation |
| M30 | 61 |  | NA |

Altogether, this points to the function and time of expression of the mentioned modules. M18 most likely clusters genes expressed before *HvFT1* as a response to hormones. Genes within M29, as *HvVRN2b*, compete with M18’s action, and do not allow the vegetative to reproductive transition. When there is enough vernalization, M20 genes are activated, repressing M29 function and the transition to reproductive phase will be effective. As the transition is already on, M18 functions are not necessary and those genes are deactivated.

### The flowering/vernalization gene regulatory network reveals differences in the interactions of *HvVRN2* genes

Transcription factors (TF) are major regulators of the floral transition, therefore to identify downstream target genes we identified a total of 2466 genes, from the BaRT2v18 catalogue, as TF (Table S8). After filtering, 1806 were part of the expression data set, and were used for gene regulatory network (GRN) construction. A GRN indicates the probability of a target gene being regulated by a TF. The GRN identified the interactions between several TF and their target genes, including well-known regulations (Table S9).

As this study focuses on vernalization, GRN was filtered to include all genic interactions of *HvVRN1*, both *HvVRN2* genes, and *HvFT1* with other TF and FT genes (Fig. 5). Twelve TF were connected by both *HvVRN1* and at least one of the *HvVRN2*, and also to *HvFT1*. The resulting network displays the genes that are most likely involved in the vernalization and flowering pathways. These genes were mostly from three co-expression modules: M18, M20 and M29. Some already known vernalization– and flowering-related genes were identified as highly related to the vernalization genes, such as both paralogs of *HvVRN1* (*HvBM3* and *HvBM8*), the circadian clock gene *HvPRR37* (*PPD-H1*), three CBFs or *HvVRT2*, a Vegetative to Reproductive Transition gene. The inclusion of these genes indicates that the experiment and methodology of analysis followed are able to identify well-known relationships. Moreover, some new transcription factors appearing in the network were also identified as differentially expressed only in C01, such as an AP2/B3 (BaRT2v18chr4HG215200), a bHLH (BaRT2v18chr6HG305650), a BZIP77-like (BaRT2v18chr5HG259930), a B3 domain-containing protein (BaRT2v18chr4HG206880), an Auxin-responsive protein (BaRT2v18chr1HG013580) or a GATA transcription factor (BaRT2v18chr4HG219040). The connections between these genes, their co-expression modules and which experimental factors correlate best with the modules allowed us to propose a temporal sequence: genes from M18 and M29 are expressed when plants are not fully vernalized, while M20 genes are activated when plants do not require vernalization or are fully vernalized. M18 and M29 regulate the transition of flowering, most likely M18 being an activator and M29 a repressor. M20 includes inducers of the transition to flowering.

**Figure 5.**
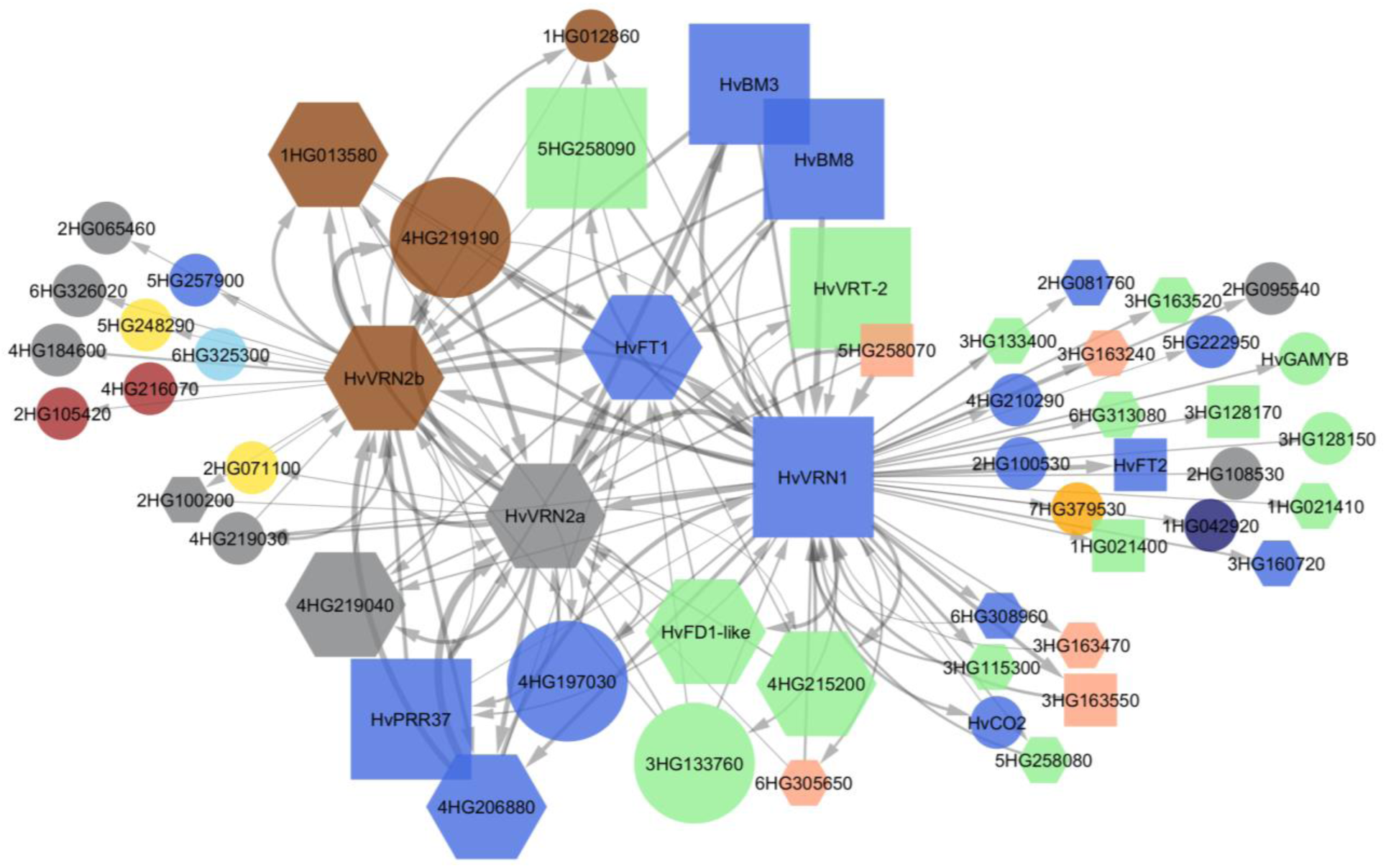
The gene regulatory network (GRN) of the three main vernalization genes *HvVRN1*, *HvVRN2* and *HvFT1* (*HvVRN3*) is similar to models proposed in the literature. The GRN shows only links between transcription factors and those three main genes. Larger node size indicates genes directly connected to *HvFT1*. Node shape determines if the gene was identified as differentially expressed due to vernalization (q-val ≤ 0.01) in both Near-Isogenic Lines (square), only in C01 genotype, with presence of *HvVRN2* (hexagon), or was not differentially expressed (circle). Node colour is assigned by co-expression module: blue for M20, green for M18, brown for M29, salmon for M27, lightblue for M28, darkblue for M15, yellow for M4, red for M3. Grey colour corresponds to M0, which includes not clustered genes. Larger link width corresponds to higher weight computed by GENIE3. Genes are labelled by their gene name or their gene ID model from BaRT2v18, trimming “BaRT2v18chr” to shorten its length, i.e. 4HG219190 as BaRT2v18chr4HG219190.

We have found genes apparently co-regulated with *HvVRN2b* (M29), and others acting in close association. One of them, *SNF2* (*helicase*), sits next to the *HvVRN2 locus* on chromosome 4H (Fig. S3). As the alleles of the two NIL genotypes at this gene are different, the CSIRO01 Hv*SNF2* allele was likely introgressed during the introduction of *HvVRN2*, by linkage drag. The other two genes in the network in the M29 module, an Auxin-responsive protein (BaRT2v18chr1HG013580) and a Histone-lysine methyltransferase (BaRT2v18chr1HG012860) are located on chromosome 1H, which is free of off-target introgressions according to 50k markers.

The network revealed interactions between *HvVRN1* and other genes, differentially expressed by vernalization, affecting *HvVRN2a* expression, but not *HvVRN2b*. This refers to transcription factors *AP2/B3*, *HvVRT2*, *HvFD1-like*, *CBF* and *bHLH*. In addition, *HvVRN2b* affects the expression of other unrelated genes. It is worth mentioning that the largest number of interactions detected were for the MADS-box gene *HvVRN1*.

### Modelling of CCT/NF-Y complexes predicted higher stability of VRN2/NF*-*Y complexes

Genes with CONSTANS-like (CCT) protein domain are heavily implicated in flowering regulation, integrating the photoperiod (*HvCO1*/*HvCO2* and *HvPRR37*) and vernalization (*HvVRN2*) pathways to control the transition to reproductive phase. Protein-protein interactions between the CCT genes have been identified *in vitro* and *in vivo* in other species, as well as with Nuclear Factor Y (NF-Y) subunits, required to trigger the expression of the florigen signal from *FT*. Still, the regulation at protein-level had not been studied in barley. We conducted an *ex situ* approach through protein modelling prediction to check if the protein-protein interactions were plausible. We investigated the three-dimensional structure of the complexes with CCT domain proteins, using AlphaFold3. Based on the models constructed for rice Hd1 and Arabidopsis CO proteins (Shen et al 2020, Lv et al 2021), the DNA-binding domain was the CCT domain rather than the zinc-finger domains of *HvCO1*/*HvCO2* or both *HvVRN2*.

We tested the stability of macrocomplexes formed between barley CCT domain containing proteins HvVRN2a, HvCO1, HvCO2 and HvPRR37 (PPD1), together with Nuclear Factor Y subunits (NF-YB and NF-YC) proteins, to bind to the DNA motif CCACA. The modelled barley CCT proteins bound fittingly to the complex containing NF-YB and NF-YC, as confirmed in rice. Overall, the complexes were more stable with HvVRN2a than with HvCO2 or HvCO1 (Fig. S5). The maximum scores (iPTM + PTM) for each of these three CCT protein with a NF-YB/NF-YC complex were 1.51, 1.37 and 1.37, respectively. Protein PPD1 also bound to the NF-Y complex, yet with lower stability (maximum score 1.3). The protein sequence of HvVRN2a was used in the complexes, since its ortholog in wheat ZCCT1 was used by Li et al (2011) in the yeast two– and three-hybrid assays. Similar results were obtained for HvVRN2b with the top 10 combinations of HvVRN2a/NF-YB/NF-YC (Table S10).

Predictions for dimers of all CCT domain proteins, either homo or heterodimers, were also tested. In all cases, they presented low-stability.

## Discussion

### Different patterns of expression between *HvVRN2* genes

This is the first time that the expression level of the two *HvVRN2* genes, over a wide range of conditions, has been clearly established. Previous reports (Trevaskis et al 2006; Szűcs et al 2007) suggested a predominant role of *HvVRN2b* over *HvVRN2a* as the main repressor in the vernalization pathway. In wheat, although both genes appear to be involved in vernalization (Distelfeld et al 2009), *ZCCT2* (ortholog of *HvVRN2b*) has a larger role (Chen and Dubcovsky 2012). Our result corroborates this hypothesis, as both genes produce essentially the same protein, and the transcripts of *HvVRN2b* are much more abundant.

According to the standing hypothesis, formulated in wheat (Chen and Dubcovsky 2012), *HvVRN1* is the repressor of *HvVRN2*. This hypothesis explains well the complete repression of *HvVRN2a* when *HvVRN1* rises. However, it is not sufficient to explain the behavior of *HvVRN2b*. Even after 60 days of vernalization, and with *HvVRN1* apparently fully expressed, some expression of *HvVRN2b* was still present. Moreover, *HvVRN2a* showed some expression at 60 days of vernalization after 25 days in the growth chamber, while it was fully repressed at 5 and 15 days, hinting at a possible de-repression (Fig. 3). Furthermore, we indicate possible TFs involved in differential expression of the two *HvVRN2* genes. We cannot conclude that one *HvVRN2* gene is more relevant than the other, but the differences in expression levels and patterns encourage further investigation for potential differences in function and environmental control. Cha et al (2022) also found strikingly different expression of the *ZCCT1* and *ZCCT2* genes in wheat, leading them to state that “…considering the *VRN2* locus as a single gene is potentially misleading in terms of understanding the vernalization response…”. Our findings are consistent with this view. The low but sustained expression of *HvVRN2b* is consistent with other reports of expression of *HvVRN2* under non-inductive conditions (Monteagudo et al 2019, Fernández-Calleja et al 2022). The protracted expression of *HvVRN2b* concurrent with *HvVRN1* and *HvFT1* full expression, suggests possible roles of *HvVRN2* beyond repression of the florigen signal.

### The computed gene regulatory network reflects known interactions and identifies new determinants of flowering

The inclusion of flowering-related genes provides a positive control for the relationships detected in our experiment. We found close relationship of *HvVRN1* expression with *HvVRN2a* and *HvVRN2b*, *HvBM3* and *HvBM8* (paralogs of *HvVRN1*), *HvFT1*, *HvFT2*, and *HvPRR37* (*PPD-H1*). Also, both *HvVRN2* genes were connected with *HvPRR37*. All these connections were already described in the literature (Mulki and von Korff 2016; Shaw et al 2020). Their detection in our experiment validates the procedures of analysis, and provides a certain confidence over the rest of the findings highlighted in Fig. 5. New connections have also been found in this study, as is the case of a GATA transcription factor or a B3 domain-containing protein. It is surprising, however, that gene expression in leaves detects relationship of genes whose expression and function occur mostly in apex. This could be an effect of the dominant allele at *PPD-H1* and, therefore, the genetic background (Digel et al 2015) on gene expression. The absence of some relationships is also informative. For instance, *HvVRN2* and *HvCO2* expression were not directly linked in our study, which is consistent with the hypothesis indicating that they compete at protein-level to bind NF-Y proteins (Li et al 2011). Moreover, *CONSTANS* expression in *Arabidopsis* is not directly linked with protein quantity. CO protein is stable in late hours under long-day photoperiod, as it is degraded during morning and night (Brambilla and Fornara 2017).

As mentioned, *HvSNF2* and *HvVRN2a* are adjacent on the long arm of chromosome 4H. Gene *HvSNF2* has a single copy in barley (Yan et al 2002). This gene seems to perform essential tasks, related to chromatin remodelling during cell division. A regulatory role in barley spike development has been suggested (Chen et al 2023), although its expression has been reported in root, spike and leaves. In this last case, higher expression was linked to cold treatments. Our study is consistent with these characteristics. Interestingly, *HvSNF2* is only expressed in CSIRO01, implying the necessity of the presence of *HvVRN2* to be activated, at least in leaves. It must be noted that *HvSNF2* and *HvVRN2a* show divergent orientations (Fig. S3). This structure could result in simultaneous expression either by the action of a common promoter region between the starting points of both genes, or by the opening of chromatin for the entire region. Another possibility could be due to the allelic differences between both genotype at this gene. Alternatively, *HvSNF2* expression could be affected by the same factors as *HvVRN2*.

*HvVRN1* plays a central role in the gene regulatory network, interacting with the largest number of genes. Our transcriptome results also support previous findings of genes controlled by VRN1, based on chromatin immunoprecipitation (Deng et al 2015). *HvFT1*, *HvVRN2*, a *bHLH* transcription factor, the zinc-finger *HvPROG1*, *CBF*s, an *AP2/B3* transcription factor, the MADS-box *HvVRT2*, and the identified novel bZIP *HvFD1-like* gene were in common with that study.

There are differences among the genes located one degree of separation from *HvVRN2a* and *HvVRN2b*, suggesting different pathways of regulation. While *HvVRN2a* seems closer to *HvFT1* and *HvVRN1*, *HvVRN2b* showed more connections to TF not related to the flowering and vernalization pathway. For example, it seems to regulate two TF within the module M3, enriched with GO related to chloroplasts and photosystems. Therefore, *HvVRN2b* could have a function beyond the flowering and vernalization pathway, as its expression continues even after the florigen signal of *HvFT1* is activated.

### Vernalization-related co-expression modules identify new candidate genes for flowering and tiller number

High correlation coefficients of modules of co-expression with vernalization suggest a close relationship of the member genes. Modules M18 and M27 show the strongest negative correlation, indicating decreasing expression as vernalization and plant age increase. The relationship of some of these genes with the vernalization pathway has already been described (Kane et al 2005; Stockinger et al 2007; Trevaskis et al 2007; Dhillon et al 2010, Li et al 2021). The MADS-box *VRT2* and *CBF* genes, associated with freezing tolerance, are expressed before *HvVRN1* is induced. After vernalization is complete, and the plant transitions to the reproductive phase, those genes are repressed.

Our research has unravelled new genes with possible function in the barley vernalization pathway. One of them is BaRT2v18chr5HG259930, ortholog of *Oryza sativa*’s BZIP77 (*OsFD1*). It is present in a module clearly affected by the application of vernalization (M18, together with *CBF* genes and *HvVRT2*). FD is the protein that forms a complex with FT1 and 14-3-3 proteins to transduce the flowering signal from the leaves to the shoot apex, as shown in *Arabidopsis* (Abe et al 2005; Wigge et al 2005) and rice (Taoka et al 2011). We have detected this gene expression related to those of *HvVRN1*, *HvVRN2a* and *HvFT1*, indicating a close relationship with the vernalization response. Li et al (2015) analysed factorial combinations of FT, FD-like and 14-3-3 proteins, promoting flowering in barley and wheat. They identified different FD-like genes but none of them corresponded to the one reported in this study.

Another interesting gene identified in this study is a zinc-finger protein, ortholog of rice *PROG1*, a gene related to leaf angle and tiller number predominantly expressed in axillary meristems (Jin et al 2008; Tan et al 2008). *OsPROG1* shows several orthologous genes in barley, all located in tandem repeats. Two *HvPROG1* genes were detected as DEGs, and were clustered by co-expression in modules M18 and M27, both negatively correlated to vernalization time. *HvPROG1* genes are annotated as zinc-finger TF but our analyses did not classify them as TF, and were not considered as regulators in the GRN analysis. Still, the two *HvPROG1* are regulated both by *HvVRN1* and *HvVRN2* (Table S9), as well as other TFs shown in Fig. 5. Karsai et al (2006), in a facultative x winter barley population, showed that tiller production is higher in presence of *HvVRN2*. In this research we have also identified differences in tiller number per day, and at Z31 and Z49 (Fig. S2). Altogether, *HvVRN2* could be an enhancer of the *HvPROG1* genes in the basal section of the leaf promoting tillering.

### Higher stability of VRN2/NF-Y complexes as limiting CO2/NF-Y activation of the flowering process

Brambilla and Fornara (2017), discussing on the combinatorial properties of the NF-Y complex, hypothesized that it could interact not only with NF-YA or CO-like proteins, but also with other CCT domain proteins in general. This has already been proven for rice GHD7-CCT, GHD8 (also named OsNF-YB11) and OsNFYC7 (Shen et al 2020).

These observations are compatible with the competition between CO2 and VRN2 described in wheat (Li et al 2011, Shaw et al 2020), and the possible differences in function of each complex. All these trimers can bind DNA through the CCT domain, and also form larger complexes among themselves, binding through their zinc-fingers. The structural analysis indicates that the zinc-finger is not the likely point of union to DNA. Rather, it is more likely that the CCT domain is the DNA binding domain, recognizing the CCACA motif present in promoter regions (Shen et al 2020).

Our results indicate that the complex of NF-Y subunits with VRN2 is more stable than with CO2, although both complexes probably co-exist and can bind to the promoter region of target genes, including *HvFT1*. Their dynamics may be affected by the strength of the larger oligomers formed by them. Zeng et al (2022) proposed that the two B-box (zinc-finger) domains of CO form a continuous head-to-tail oligomer-like structure, further stabilized by the binding of the CO-CCT/NF-Y complexes, through the CCT domains to the *FT1* promoter, inducing its expression. This working model was experimentally confirmed in *Arabidopsis thaliana* (Huang et al 2025). In barley, the stability of this complex structure could be affected by the lesser number of zinc-finger domains carried by VRN2 (only one) and PPD1 (none), not allowing the head-to-tail binding structure, hence affecting *HvFT1* expression. VRN2 could compete with CO2 to bind NF-Y and, additionally the presence of VRN2/NF-Y concurrently with CO2/NF-Y could weaken the stability of the protein complexes binding to the *HvFT1* promoter. Moreover, the differences in transcript quantity between *HvVRN2* genes could be explained due to the extra CCACA DNA motif present in the promoter of *HvVRN2b*, compared to *HvVRN2a* (Fig. S6). This extra binding-site could allow a more stable oligomerization of the CCT/NF-Y complex, explaining the differences in expression between the *HvVRN2* genes (Fig. 2).

Summarizing, differences in expression of both *HvVRN2* genes have been identified. *HvVRN2a* gene expression is lower than that of *HvVRN2b*. *HvVRN2b* is still expressed in early reproductive phase and *HvVRN2a* expression slightly increased in plants fully vernalized, after 25 days under long-day photoperiod. While both *HvVRN2* genes seem to regulate the flowering pathway, a gene regulatory network identified connections outside the florigen signal of *HvFT1*, mostly for *HvVRN2b*. While both *HvVRN2* have a redundant repressor effect in flowering, they should be considered as different genes. The expression of *HvVRN2* locus increases during the accumulation of days under long-day photoperiod, after insufficient vernalization, suggesting a possible interaction with *HvPRR37 (PPD-H1)*. While CCT proteins (including VRN2) compete for the NF-Y complex, VRN2/NF-Y complexes were predicted as more stable. The higher stability of VRN2/NF-Y complexes could block the CO2/NF-Y oligomerization required to induce *HvFT1*. Moreover, the higher tillering of winter barley could be related to *HvPROG1* genes, induced by *HvVRN2*.

In conclusion, the *ZCCT* genes inside locus *HvVRN2* should be considered as different genes and could have different potential roles during vernalization. The repression of flowering by locus *HvVRN2* may take place at protein level, more likely preventing the required oligomerization of CO2/NF-YB/NF-YC protein complexes at the promoter region of *HvFT1*. *HvVRN2* has a potential role in tiller production, which could translate into higher yields. Further research on *HvVRN2* should consider the differences between the *ZCCT* genes.

## Supplementary data

Supplementary Table S1. Sample information and sequencing data

Supplementary Table S2. Expression (counts and tpm) at gene-level for all samples

Supplementary Table S3. Flowering-related gene names and gene model ID in BaRT2v18 transcriptome

Supplementary Table S4. Protein sequences and oligonucleotide with CCACA DNA-motif for modelling prediction

Supplementary Table S5. Differentially expressed genes by vernalization for both genotypes, separately

Supplementary Table S6. Co-expression module membership of expressed genes

Supplementary Table S7. Gene Ontology enrichment for each co-expression module

Supplementary Table S8. Identified transcription factors in the reference transcriptome

Supplementary Table S9. Gene Regulatory Network, in pairwise relations of regulator and its target

Supplementary Table S10. Modelling scores of CCT/CCT dimers and CCT/NF-Y protein complexes

Supplementary Figure S1. Photos of unvernalized plants, showing differences in development

Supplementary Figure S2. Tiller production per day, and at Z31 and Z49

Supplementary Figure S3. Pangenome occupancy of *HvSNF2* and *HvVRN2*

Supplementary Figure S4. Scatter plot of CCT/NF-Y modelled protein complexes and 3D structure of VRN2a/NF-Y complex

Supplementary Figure S5. Location of CCACA motif in the promoters of CCT genes

## Acknowledgements

We thank Ben Trevaskis’ group for the near-isogenic lines developed at CSIRO Agriculture and Food, Canberra, Australia (GRDC-funded project, CSP00183). FMT thanks Asún Costar and Vanesa Martínez Agudo for their help in plant growth.

## Author contributions

Funding was acquired by AMC, BCM, EI, FMT and IK. The experiment was designed by AMC, EI, FMT and IK. Plant growth and wet-lab was performed by AMC, FMT and IK. Dry-lab methodology was designed by BCM, FMT and PB. Data was analysed by FMT and IP. Research was supervised by AMC, BCM, EI, IK and PB. Summarized data and graphics were created by AMC, BCM, FMT, IP and EI. Original draft was prepared by AMC, FMT and EI. All authors revised and approved the final manuscript.

## Bibliography

1. Abe M, Kobayashi Y, Yamamoto S, Daimon Y, Yamaguchi A, Ikeda Y, Ichinoki H, Notaguchi M, Goto K, Araki T. (2005). FD, a bZIP protein mediating signals from the floral pathway integrator FT at the shoot apex. Science 309, 1052–1056

2. Abramson J, Adler J, Dunger J, Evans R, Green T, Pritzel A, Ronneberger O, Willmore L, Ballard AJ, Bambrick J, Bodenstein SW, Evans DA, Hung CC, O’Neill M, Reiman D, Tunyasuvunakool K, Wu Z, Žemgulytė A, Arvaniti E, Beattie C, Bertolli O, Bridgland A, Cherepanov A, Congreve M, Cowen-Rivers AI, Cowie A, Figurnov M, Fuchs FB, Gladman H, Jain R, Khan YA, Low CMR, Perlin K, Potapenko A, Savy P, Singh S, Stecula A, Thillaisundaram A, Tong C, Yakneen S, Zhong ED, Zielinski M, Žídek A, Bapst V, Kohli P, Jaderberg M, Hassabis D, Jumper JM (2024). Accurate structure prediction of bio-molecular interactions with AlphaFold 3. Nature 630, 493–500

3. Brambilla V, Fornara F. (2017). Y flowering? Regulation and activity of CONSTANS and CCT-domain proteins in Arabidopsis and crop species. Biochim. Biophys. Acta 1860, 655–660

4. Bray N, Pimentel H, Melsted P, Pachter L (2016). Near-optimal probabilistic RNA-seq quantification. Nat. Biotechnol. 34, 525–527

5. Cantalapiedra CP, García-Pereira MJ, Gracia MP, Igartua E, Casas AM, Contreras-Moreira B. (2017). Large differences in gene expression responses to drought and heat stress between elite barley cul-tivar Scarlett and Spanish landrace. Front. Plant Sci. 8:647

6. Carrera SC, Savin R, Slafer GA (2024). Critical period for yield determination across grain crops. Trends Plant Sci. 29, 329–342

7. Cha JK, O’Connor K, Alahmad S, Lee JH, Dinglasan E, Park H, Lee SM, Hirsz D, Kwon SW, Kwon Y, Kim YM, Ko JM, Hickey LT, Shin D, Dixon LE. (2022). Speed vernalization to accelerate generation advance in winter cereal crops. Mol. Plant. 15, 1300–1309.

8. Chen A, Dubcovsky J. (2012). Wheat TILLING mutants show that the vernalization gene *VRN1* down-regulates the flowering repressor *VRN2* in leaves but is not essential for flowering. PLoS Genet 8:e1003134

9. Chen S, Zhou Y, Gu J. (2018). fastp: an ultra-fast all-in-one FASTQ preprocessor. Bioinformatics 34, i884–i890

10. Chen G, Mishina K, Zhu H, Kikuchi S, Sassa H, Oono Y, Komatsuda T. (2023). Genome-wide analysis of Snf2 gene family reveals potential role in regulation of spike development in barley. Int. J. Mol. Sci. 24:457

11. Contreras-Moreira B, Saraf S, Naamati G, Casas AM, Amberkar SS, Flicek P, Jones AR, Syer S. (2023). GET_PANGENES: calling pangenes from plant genome alignments confirms presence-absence vari-ation. Genome Biol. 24:223

12. Corbesier L, Vincent C, Jang S, Fornara F, Fan Q, Searle I, Giakountis A, Farrona S, Gissot L, Turnbull C, Coupland G. (2007). FT protein movement contributes to long-distance signaling in floral induc-tion of *Arabidopsis*. Science 316, 1030–1033

13. Coulter M, Entizne JC, Guo W, Bayer M, Wonneberger R, Milne L, Schreiber M, Haaning A, Mueh-lbauer G, McCallum N, Fuller J, Simpson C, Stein N, Brown JWS, Waugh R, Zhang R. (2022). BaRTv2: a highly resolved barley reference transcriptome for accurate transcript-specific RNA-seq quantifi-cation. Plant J. 111, 1183–1202

14. Deng W, Casao MC, Wang P, Sato K, Hayes PM, Finnegan EJ, Trevaskis B. (2015). Direct links between the vernalization response and other key traits of cereals crops. Nat. Commun. 6:5882.

15. Dhillon T, Pearce SP, Stockinger EJ, Distelfeld A, Li C, Knox AK, Vashegyi I, Vágújfalvi A, Galiba G, Dubcovsky J. (2010). Regulation of freezing tolerance and flowering in temperate cereals: The *VRN-1* connection. Plant Physiol. 153, 1846–1858

16. Digel B, Pankin A, von Korff M. (2015). Global transcriptome profiling of developing leaf and shoot apices reveals distinct genetic and environmental control of floral transition and inflorescence de-velopment in barley. Plant Cell 27, 2318–2334

17. Distelfeld A, Tranquilli G, Li C, Yan L, Dubcovsky J. (2009). Genetic and molecular characterization of the VRN2 loci in tetraploid wheat. Plant Physiol. 149, 245–257

18. Dubcovsky J, Chen C, Yan L. (2005). Molecular characterization of the allelic variation at the VRN-H2 vernalization locus in barley. Mol. Breeding 15, 395–407

19. Fernández-Calleja M, Casas AM, Igartua E. (2021). Major flowering time genes of barley: allelic di-versity, effects, and comparison with wheat. Theor. Appl. Genet. 134, 1867–1897

20. Fernández-Calleja M, Ciudad FJ, Casas AM, Igartua E. (2022). Hybrids provide more options for fine-tuning flowering time responses of winter barley. Front. Plant Sci. 13:827701.

21. Gnesutta N, Kumimoto RW, Swain S, Chiara M, Siriwardana C, Horner DS, Holt III BF, Mantovani R. (2017). CONSTANS imparts DNA sequence specificity to the histoen fold NF-YB/NF-YC dimer. Plant Cell. 29, 1516–1532

22. Homma F, Huang J, van der Hoorn RAL (2023). AlphaFold-Multimer predicts cross-kingdom interac-tions at the plant-pathogen interface. Nat. Commun. 14:6040

23. Huang X, Ma Z, He D, Han X, Liu X, Dong Q, Tan C, Yu B, Sun T, Nordenskiöld L, Lu L, Miao Y, Hou X (2025). Molecular condensation of the CO/NF-YB/NF-YC/FT complex gates floral transition in *Ara-bidopsis*. EMBO J. 44, 225–250

24. Huynh-Thu V, Irrthum A, Wehenkel L, Geurts P. (2010). Inferring regulatory networks from expres-sion data using tree-based methods. PLoS ONE 5:e12776

25. Jayakodi M, Padmarasu S, Haberer G, Bonthala VS, Gundlach H, Monat C, Lux T, Kamal N, Lang D, Himmelbach A, Ens J, Zhang XQ, Angessa TT, Zhou G, Tan C, Hill C, Wang P, Schreiber M, Boston LB, Plott C, Jenkins J, Guo Y, Fiebig A, Budak H, Xu D, Zhang J, Wang C, Grimwood J, Schmutz J, Guo G, Zhang G., Mochida K, Hirayama T, Sato K, Chalmers KJ, Langridge P, Waugh R, Pozniak CJ, Scholz Y, Mayer KFX, Spannagl M, Li C, Mascher M, Stein N. (2020). The barley pan-genome reveals the hidden legacy of mutation breeding. Nature 588, 284–289

26. Jin J, Huang W, Gao JP, Yang J, Shi M, Zhu MZ, Luo D, Lin HX. (2008). Genetic control of rice plant architecture under domestication. Nat. Genet. 40, 1365–1369

27. Kane NA, Danyluk J, Tardif G, Ouellet F, Laliberté JF, Limin AE, Fowler DB, Sarhan F. (2005). TaVRT-2, a member of the St MADS-11 clade of flowering repressors, is regulated by vernalization and photoperiod in wheat. Plant Physiol. 138, 2354–2363

28. Karsai I, Mészáros K, Szűcs P, Hayes PM, Láng L, Bedő Z (2006). The influence of photoperiod on the *Vrn-H2* locus (4H) which is a major determinant of plant development and reproductive fitness traits in a facultative *x* winter barley (*Hordeum vulgare* L.) mapping population. Plant Breed. 125, 468–472

29. Kovacik M, Nowicka A, Zwyrtková J, Strejčková B, Vardanega I, Esteban E, Pasha A, Kaduchová K, Krautsova M, Červenková M, Šafář J, Provart NJ, Simon R, Pecinka A. (2024). The transcriptome land-scape of developing barley seeds. Plant Cell 36, 2512–2530

30. Langfelder P, Horvath S. (2008). WGCNA: an R package for weighted correlation network analysis. BMC Bionformatics 9:559

31. Li W, Godzik A. (2006). Cd-hit: a fast program for clustering and comparing large sets of protein or nucleotide sequences. Bioinform. 22, 1658–1659

32. Li C, Distelfeld A, Comis A, Dubcovsky J. (2011). Wheat flowering repressor VRN2 and promoter CO2 compete for interactions with NUCLEAR FACTOR-Y complexes. Plant J. 67, 762–773.

33. Li C, Lin H, Dubcovsky J. (2015). Factorial combinations of protein interactions generate a multiplicity of florigen activation complexes in wheat and barley. Plant J. 84, 70–82.

34. Li C, Lin H, Chen A, Lau M, Jernstedt J, Dubcovsky J. (2019). Wheat VRN1, FUL2 and FUL3 play critical and redundant roles in spikelet development and spike determinacy. Development 146, dev175398

35. Li K, Debernardi JM, Li C, Lin H, Zhang C, Jernstedt J, von Korff M, Zhong J, Dubcovsky J. (2021). Interactions between SQUAMOSA and SHORT VEGETATIVE PHASE MADS-box proteins regulate me-ristem transitions during wheat spike development. Plant Cell 33, 3621–3644

36. Li T, Li Y, Shangguan H, Bian J, Luo R, Tian Y, Li Z, Nie X, Cui L. (2023). BarleyExpDB: an integrative gene expression database for barley. BMC Plant Biol. 23:170

37. Love MI, Huber W, Anders S. (2014). Moderated estimation of fold change and dispersion for RNA-seq data with DESeq2. Genome Biol. 15:550

38. Lv Xinchen, Zeng X, Hu H, Chen L, Zhang F, Liu R, Liu Y, Zhou X, Wang C, Wu Z, Kim C, He Y, Du J. (2021). Structural insights into the multivalent binding of the Arabidopsis *FLOWERING LOCUS T* pro-moter by the CO-NF-Y master transcription factor complex. Plant Cell 33, 1182–1195

39. Mascher M, Wicker T, Jenkins J, Plott C, Lux T, Koh CS, Ens J, Gundlach H, Boston LB, Tulpová Z, Holden S, Hernández-Pinzón I, Scholz W, Mayer KFX, Spannagl M, Pozniak CJ, Sharpe AG, Šimková H, Moscou MJ, Grimwood J, Schmutz J, Stein N. (2021). Long-read sequence assembly: a technical evaluation in barley. Plant Cell 33, 1888–1906

40. Milne L, Bayer M, Rapazote-Flores P, Mayer CD, Waugh R, Simpson CG. (2021). EORNA, a barley gene and transcript abundance database. Sci. Data 8:90

41. Monteagudo A, Igartua E, Contreras-Moreira B, Gracia MP, Ramos J, Karsai I, Casas AM. (2019). Fine-tuning of the flowering time control in winter barley: the importance of *HvOS2* and *HvVRN2* in non-inductive conditions. BMC Plant Biol. 19:113

42. Mulki MA, von Korff M. (2016). *CONSTANS* controls floral repression by up-regulating *VERNALIZA-TION2* (*VRN-H2*) in barley. Plant Physiol. 170, 325–337

43. Müller LM, Mombaerts L, Pankin A, Davis SJ, Webb AAR, Goncalves J, von Korff M. (2020). Differen-tial effects of day/night cues and the circadian clock on the barley transcriptome. Plant Physiol. 183, 765–779

44. Ochagavía H, Kiss T, Karsai I, Casas AM, Igartua E. (2022). Responses of barley to high ambient tem-perature are modulated by vernalization. Front. Plant Sci. 12:776982

45. Panahi B, Mohammadi SA, Ruzicka K, Holaso HA, Mehrjerdi MZ. (2019). Genome-wide identification and co-expression network analysis of nuclear factor-Y in barley revealed potential functions in salt stress. Physiol Mol Biol Plants 25, 485–95

46. Pimentel H, Bray N, Puente S, Melsted P, Pachter L. (2017). Differential analysis of RNA-seq incorpo-rating quantification uncertainty. Nat. Methods 14, 687–690

47. R Core Team (2023). R: A language and environment for statistical computing

48. Rapazote-Flores P, Bayer M, Milne L, Mayer CD, Fuller J, Guo W, Hedley PE, Morris J, Halpin C, Kam J, McKim SM, Zwirek M, Casao MC, Barakate A, Schreiber M, Stephen G, Zhang R, Brown JWS, Waugh R, Simpson CG. (2019). BaRTv1.0: an improved barley reference transcript dataset to determine ac-curate changes in the barley transcriptome using RNA-seq. BMC Genomics 20:968

49. Santana-Garcia W, Castro-Mondragon JA, Padilla-Gálvez M, Nguyen NTT, Elizondo-Salas A, Ksouri N, Gerbes F, Thieffry D, Vincens P, Contreras-Moreira B, van Helden J, Thomas-Collier M, Medina-Rivera A. (2022). RSAT 2022: regulatory sequence analysis tools. Nucleic Acids Res. 50, 670–676

50. Sasani S, Hemming MN, Oliver SN, Greenup A, Mahfoozi S, Poustini K, Sharifi H, Dennis ES, Peacock WJ, Trevaskis B. (2009). The influence of vernalization and daylength on expression of flowering-time genes in the shoot apex and leaves of barley (*Hordeum vulgare*). J. Exp. Bot. 60, 2169–2178

51. Schloerke B, Cook D, Larmarange J, Briatte F, Marbach M, Thoen E, Elberg A, Crowley J. (2024). Ggally: extension to “ggplot2”. R package version 2.2.1, https://github.com/ggobi/ggally, https://ggobi.github.io/ggally/.

52. Shannon P, Markiel A, Ozier O, Baliga NS, Wang JT, Ramage D, Amin N, Schwikowski B, Ideker T. (2003). Cytoscape: a software environment for integrated models of biomolecular interaction net-works. Genome Res. 13, 2498–2504

53. Shaw LM, Li C, Woods DP, Alvarez MA, Lin H, Lau MY, Chen A, Dubcovksy J. (2020). Epistatic inter-actions between PHOTOPERIOD1, CONSTANS1 and CONSTANS2 modulate the photoperiodic re-sponse in wheat. PLoS Genet 16:e1008812

54. Shen C, Liu H, Guan Z, Yan J, Zheng T, Yan W, Wu C, Zhang Q, Yin P, Xing Y. (2020). Structural insight into DNA recognition by CCT/NF-YB/YC complexes in plant photoperiodic flowering. Plant Cell 32, 3469–3484

55. Sievers F, Higgins DG. (2014). Clustal Omega, accurate alignment of very large numbers of se-quences. Methods Mol Biol. 1079, 105–116

56. Soneson C, Love MI, Robinson MD. (2015). Differential analyses for RNA-seq: transcript-level esti-mates improve gene-level inferences. F1000Res. 4:1521

57. Stockinger EJ, Skinner JS, Gardner KG, Francia E, Pecchioni N. (2007). Expression levels of barley *Cbf* genes at the *Frost resistance-H2* locus are dependent upon alleles at *Fr-H1* and *Fr-H2*. Plant J. 51, 308–321

58. Szűcs P, Skinner JS, Karsai I, Cuesta-Marcos A, Haggard KG, Corey AE, Chen THH, Hayes PM. (2007). Validation of the *VRN-H2*/*VRN-H1* epistatic model in barley reveals that intron length variation in *VRN-H1* may account for a continuum of vernalization sensitivity. Mol. Genet. Genomics 277, 249– 261

59. Tamaki S, Matsuo S, Wong HL, Yokoi S, Shimamoto K. (2007). Hd3a protein Is a mobile flowering signal in rice. Science 316, 1033–1036

60. Tan L, Li X, Liu F, Sun X, Li C, Zhu Z, Fu Y, Cai H, Wang X, Xie D, Sun C. (2008). Control of a key transition from prostrate to erect growth in rice domestication. Nat. Genet. 40, 1360–1364

61. Taoka KI, Ohki I, Tsuji H, Furuita K, Hayashi K, Yanase T, Yamaguchi M, Nakashima C, Purwestri YA, Tamaki S, Ogaki Y, Shimada C, Nakagawa A, Kojima C, Shimamoto K. (2011). 14-3-3 proteins act as intracellular receptors for rice Hd3a florigen. Nature 476, 332–335

62. Thiel J, Koppolu R, Trautewig C, Hertig C, Kale SM, Erbe S, Mascher M, Himmelbach A, Rutten T, Esteban E, Pasha A, Kumlehn J, Provart NJ, Vanderauwera S, Frohberg C, Schnurbusch T. (2021). Transcriptional landscapes of floral meristems in barley. Sci. Adv. 7:eabf0832

63. Trevaskis B, Bagnall DV, Hellis MH, Peacock J, Dennis ES. (2003). MADS box genes control vernaliza-tion-induced flowering in cereals. Proc. Natl. Acad. Sci. USA 100, 13099–13104

64. Trevaskis B, Hemming MN, Peacock WJ, Dennis ES. (2006). HvVRN2 responds to daylength, whereas HvVRN1 is regulated by vernalization and developmental status. Plant Physiol. 140, 1397–1405

65. Trevaskis B, Tadege M, Hemming MN, Peacock WJ, Dennis ES, Sheldon C. (2007). *Short Vegetative Phase*-like MADS-box genes inhibit floral meristem identity in barley. Plant Physiol. 143, 225–235

66. Wigge PA, Kim MC, Jaeger KE, Busch W, Schmid M, Lohmann JU, Weigel D. (2005). Integration of spatial and temporal information during floral induction in Arabidopsis. Science 309, 1056–1059

67. Wu Y, Hu E, Xu S, Chen M, Guo P, Dai Z, Feng T, Zhou L, Tang W, Zhan L, Fu X, Liu S, Bo X, Yu G. (2021). clusterProfiler 4.0: A universal enrichment tool for interpreting omics data. Innovation 2:100141

68. Yan L, Echenique V, Busso C, SanMiguel P, Ramakrishna W, Bennetzen JL, Harrington S, Dubcovsky J. (2002). Cereal genes similar to *Snf2* define a new subfamily that includes human and mouse genes. Mol. Genet. Genomics 268, 488–499

69. Yan L, Loukoianov A, Tranquilli G, Helguera M, Fahima T, Dubcovsky J. (2003). Positional cloning of the wheat vernalization gene *VRN1*. Proc. Natl. Acad. Sci. USA 100, 6263–6268

70. Yan L, Loukoianov A, Blechl A, Tranquilli G, Ramakrishna W, Sanmiguel P, Bennetzen JL, Echenique V, Dubcovsky J. (2004). The wheat *VRN2* gene is a flowering repressor down-regulated by vernaliza-tion. Science 303, 1640–1644.

71. Yan L, Fu D, Li C, Blechl A, Tranquilli G, Bonafede M, Sanchez A, Valarik M, Yasuda S, Dubcovsky J. (2006). The wheat and barley vernalization gene *VRN3* is an orthologue of *FT*. Proc. Natl. Acad. Sci. USA 103, 19581–19586

72. Zadoks JC, Chang TT, Konaz CF. (1974). A decimal code for the growth stages of cereals. Weed Res. 14, 415–421

73. Zeng X, Lv X, Liu R, He H, Liang S, Chen L, Zhang F, Chen L, He Y, Du J. (2022). Molecular basis of CONSTANS oligomerization in FLOWERING LOCUS T activation. J. Integr. Plant Biol. 64, 731–740

74. Zhang H, Sreenivasulu N, Weschke W, Stein N, Rudd S, Radchuk V, Potokina E, Scholz U, Schweizer P, Zierold W, Langridge P, Varshney RK, Wobus U, Graner A. (2004). Large-scale analysis of the barley transcriptome based on expressed sequence tags. Plant J. 4, 276–290

75. Zheng Y, Jiao C, Sun H, Rosli HG, Pombo MA, Zhang P, Banf M, Dai X, Martin GB, Giovannoni JJ, Zhao PX, Rhee SY, Fei Z. (2016). iTAK: A program for genome-wide prediction and classification of plant transcription factors, transcriptional regulators, and protein kinases. Mol. Plant 9, 1667–1670

